# Extreme high-elevation mammal surveys reveal unexpectedly high upper range limits of Andean mice

**DOI:** 10.1101/2023.08.22.554215

**Authors:** Jay F. Storz, Marcial Quiroga-Carmona, Schuyler Liphardt, Naim M. Bautista, Juan C. Opazo, Adriana Rico Cernohorska, Jorge Salazar-Bravo, Jeffrey M. Good, Guillermo D’Elía

**Affiliations:** School of Biological Sciences, University of Nebraska, Lincoln, NE, United States; Instituto de Ciencias Ambientales y Evolutivas, Facultad de Ciencias, Universidad Austral de Chile, Valdivia, Chile; Colección de Mamíferos, Facultad de Ciencias, Universidad Austral de Chile, Campus Isla Teja, Valdivia, Chile; Division of Biological Sciences, University of Montana, Missoula, MT, United States; Millennium Nucleus of Ion Channels-Associated Diseases (MiNICAD); Facultad de Medicina y Ciencia, Universidad San Sebastián, Valdivia, Chile; Integrative Biology Group, Valdivia, Chile; Colecciόn Boliviana de Fauna, Instituto de Ecología, Universidad Mayor de San Andrés, La Paz, Bolivia; Department of Biological Sciences, Texas Tech University, Lubbock, TX, United States

**Keywords:** Andes, distribution limits, high altitude, *Phyllotis*, Puna de Atacama, species limits

## Abstract

In the world’s highest mountain ranges, uncertainty about the upper elevational range limits of alpine animals represents a critical knowledge gap regarding the environmental limits of life and presents a problem for detecting range shifts in response to climate change. Here we report results of mountaineering mammal surveys in the Central Andes, which led to the discovery of multiple species of mice living at extreme elevations that far surpass previously assumed range limits for mammals. We live-trapped small mammals from ecologically diverse sites spanning >6700 m of vertical relief, from the desert coast of northern Chile to the summits of the highest volcanoes in the Andes. We used molecular sequence data and whole-genome sequence data to confirm the identities of species that represent new elevational records and to test hypotheses regarding species limits. These discoveries contribute to a new appreciation of the environmental limits of vertebrate life.

## Introduction

Identifying factors that influence geographic range limits of animal species is a longstanding goal in ecology and evolutionary biology. Elevational shifts in species’ ranges are of special interest in the context of global climate change (Moritz et al. 2008; La Sorte and Jetz 2010; Rowe et al. 2010; Tingley et al. 2012; McCain et al. 2021; Storz and Scott 2023), and the conservation implications of such shifts underscore the need for accurate information about contemporary range limits. However, in the most mountainous regions of the planet – where potential elevational range limits are the highest – the upper limits of species’ ranges are often poorly demarcated due to a paucity of survey data. For example, in the South American rodent genus *Phyllotis* (Cricetidae: Sigmodontinae), latitudinal limits of nominal species are relatively well-delineated (Steppan and Ramirez 2015; Jayat et al. 2021; Ojeda et al. 2021), but the upper elevational limits of species with montane distributions are known with far less certainty. This was highlighted by results of a recent mountaineering mammal survey of Volcán Llullaillaco (Argentina-Chile) which recorded a new elevational record for *Phyllotis vaccarum* (previously recognized as *P. xanthopygus rupestris*) from the summit of the volcano at 6739 m (Storz et al. 2020). This record far surpasses previous records for *Phyllotis* and represents the highest specimen-based record for any mammalian species. This discovery highlights the need for novel survey approaches to document the elevational range limits of small mammals in remote, highly mountainous regions such as the Andean Altiplano.

Here we report results of five high-elevation mammal surveys (2020-2022) in the Puna de Atacama of northern Chile and bordering regions of Argentina and Bolivia. By conducting biological surveys at extreme elevations that had never been surveyed, we tested the hypothesis that small mammals inhabit much higher elevations than previously supposed. The high-elevation surveys of Andean mammals were designed to help redress the so-called Wallacean shortfall (Hortal et al. 2015), which refers to the discrepancy between the known distributional limits of species and their actual limits.

The Central Andean dry puna ecoregion is characterized by montane grasslands and shrublands interspersed with salt flats, high plateaus, and snow-capped volcanoes. Our high-elevation surveys focused on 21 volcanoes distributed along the spine of the Andean Cordillera and surrounding regions in the Altiplano. To obtain a broad picture of elevational distributions of small mammals in this remote and largely unexplored region, and to demarcate lower and upper range limits, we also surveyed small mammals at lower elevations in the Atacama Desert and along the Pacific coast of northern Chile. We report elevational records for multiple species, providing a new appreciation of the environmental tolerances of mammals and the habitable limits of the high mountain biome.

## Material and Methods

### Elevational Surveys

In high-elevation expeditions between 2020 and 2022, we trapped small mammals at sites that spanned a diversity of habitats over a broad range of elevations (fig. 1*A*, table S1). For each of 21 surveyed volcanoes (fig. 1*B*, table S2), we maintained a base camp for multiple days and maintained trap lines in the general vicinity, typically at elevations between 3800-5100 m. In the case of eight volcanoes with summits 6046-6893 m (Nevado Sajama, Parinacota, Aucanquilcha, Acamarachi, Púlar, Socompa, Llullaillaco, and Ojos del Salado), we also maintained trap lines for 1-4 days at especially high elevations between 5100-5850 m.

**Figure 1.**
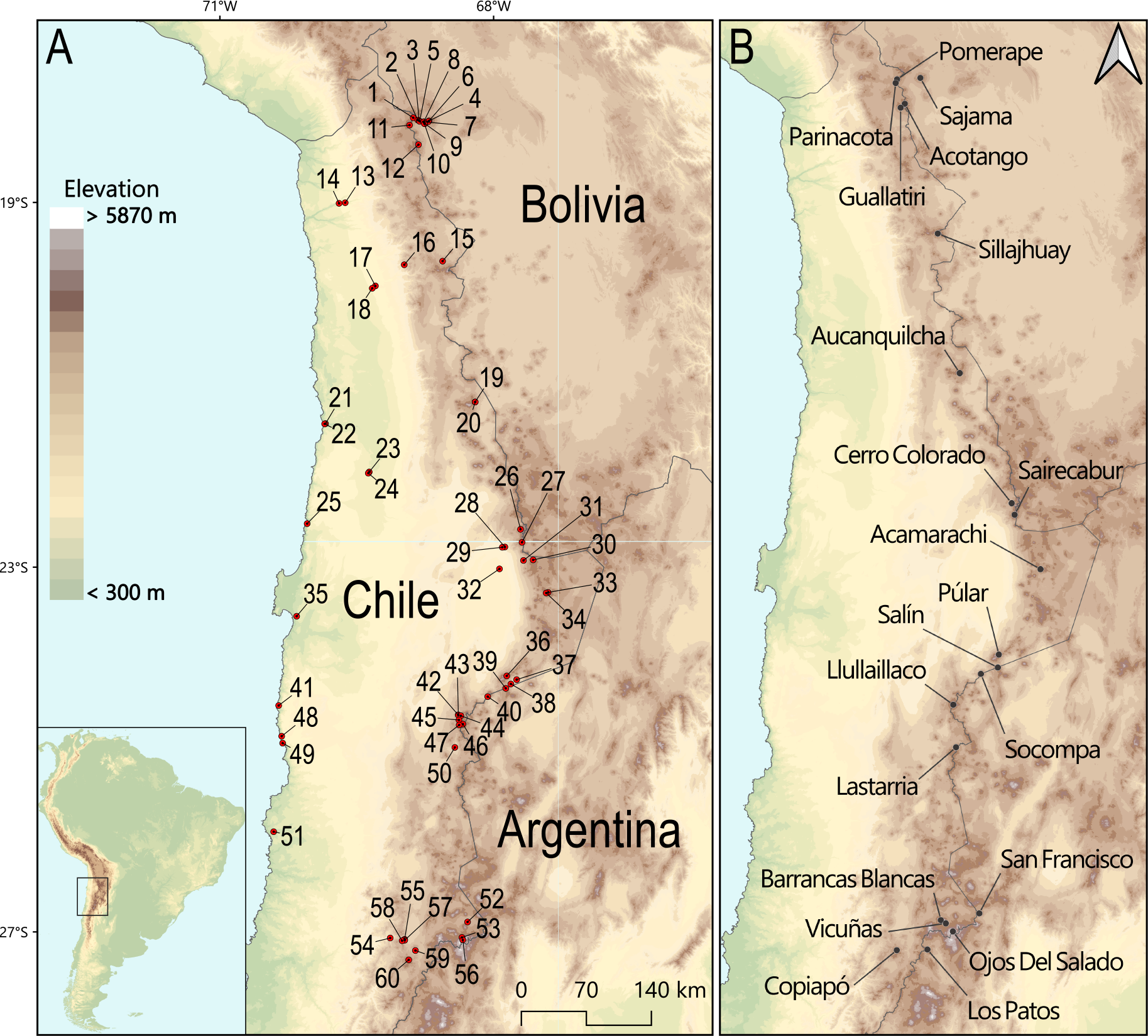
Collection localities. (A) 60 collection localities in northern Chile and bordering regions of Argentina and Bolivia, with elevations ranging from sea level to 6739 m (see table S1 for details). (B) Location of 21 surveyed volcanoes along the spine of details).

#### Live-Trapping and Specimen Preparation

We captured all mice using Sherman live traps, in combination with a smaller number of Museum Special snap traps at some sites. We sacrificed mice in the field, prepared them as museum specimens, and preserved tissue samples as sources of DNA. All specimens are housed in the Colección de Mamíferos of the Universidad Austral de Chile, Valdivia, Chile, or in the Colección Boliviana de Fauna, La Paz, Bolivia. We identified all specimens to the species level based on external characters (Patton et al. 2015) and we later confirmed all uncertain field-identifications with DNA sequence data.

In Chile, all animals were collected in accordance with permissions to JFS and GD from the following Chilean government agencies: Servicio Agrícola y Ganadero (SAG, Resolución exenta #’s 6633/2020, 2373/2021, 5799/2021, 3204/2022, and 3565/2022), Corporación Nacional Forestal (CONAF, Autorización #’s 171219 and 1501221), and Dirección Nacional de Fronteras y Límites del Estado (DIFROL, Autorización de Expedición Científica #68 and 02/22). In Bolivia, all mice were collected in accordance with permissions to ARC, JS-B, and JFS from the Ministerio de Medio Ambiente y Agua, Estado Plurinacional de Bolivia (Resolución Administrativa 026/09). All live-trapped mice were handled in accordance with protocols approved by the Institutional Animal Care and Use Committee of the University of Nebraska (project ID’s: 1919, 2100) and the bioethics committee of the Universidad Austral de Chile (certificate 456/2022).

#### Elevational vegetation surveys

For each of the 21 volcanoes that we surveyed, we treated the ascent route as an elevational transect and we recorded the upper limits of vegetation. In many cases, we traversed the same route multiple times over several consecutive days, for example, when making repeated trips from base camp to high camp.

### Molecular Sequence Data and Whole-Genome Polymorphism Data

#### DNA Sequencing

We sequenced the mitochondrial *cytochrome b* gene (*cytb*) of 82 *Phyllotis* specimens for the purpose of confirming field-based species identifications. We extracted DNA from liver samples and we PCR-amplified the first 801 base pairs of *cytb* using the primers MVZ 05 and MVZ 16 (Smith and Patton 1993), following protocols of Salazar-Bravo et al. (2001) and Cadenillas and D’Elía (2021). All newly generated sequences were deposited in GenBank (XXXXXXX-XXXXXXX).

#### Taxon sampling for analysis of whole-genome sequence data

Using a chromosome-level reference genome for *Phyllotis vaccarum* (Storz et al. 2023), we generated low-coverage whole-genome sequence data for a subset of 61 *Phyllotis* specimens to resolve uncertainty regarding species limits. We generated new WGS data for 17 *Phyllotis* specimens which we then analyzed in conjunction with previously published genome sequence data for 44 specimens (Storz et al. 2023).

#### Genomic library preparation and whole-genome sequencing

All library preparations for whole genome resequencing experiments were conducted in the University of Montana Genomics Core facility. We extracted genomic DNA from ethanol-preserved liver tissue using the DNeasy Blood and Tissue kit (Qiagen). We used a Covaris E220 sonicator to shear DNA and we then prepared genomic libraries using the KAPA HyperPlus kit (Roche). Individual libraries were indexed using KAPA UDI’s and pooled libraries were sent to Novogene for Illumina paired-end 150 bp sequencing on a Novaseq X.

#### Read quality processing and mapping to the reference genome

We used fastp 0.23.2 (Chen et al. 2018) to remove adapter sequences, and to trim and filter low-quality reads from sequences generated from library preparations. We used a 5 bp sliding window to remove bases with a mean quality less than 20 and we discarded all reads <25 bp. We merged all overlapping reads that passed filters and retained all reads that could not be merged or whose paired reads failed filtering. We separately mapped merged reads, unmerged but paired reads, and unpaired reads to the *P. vaccarum* reference genome with BWA 0.7.17 (Li and Durbin 2009) using the mem algorithm with the -M option which flags split reads as secondary for downstream compatibility. We sorted, merged, and indexed all resulting binary alignment maps with SAMtools 1.15.1 (Li et al. 2009). We used picard 2.27.4 to detect and remove PCR duplicates. We used GATK 3.8 (McKenna et al. 2010) to perform local realignment around targeted indels to generate the final BAM files.

#### Analysis of genomic variation

We calculated genotype likelihoods for scaffolds 1-19 (covering >90% of the genome) for all samples in ANGSD 0.939 (Korneliussen et al. 2014). We used -GL 1 to specify the SAmtools model for genotype likelihoods, retained only sites with a probability of being variable >1e-6 with -SNP_pval 1e-6. We filtered out bad and non-uniquely mapped reads with -remove_bads 1 and -uniqueOnly 1, respectively, and only retained reads and bases with a mapping quality higher than 20. We adjusted mapping quality for excessive mismatches with -C 50 and recalculated base alignment quality with -baq 1. We used PCAngsd v.0.99.0 (Meisner and Albrechtsen 2018) to calculate the covariance matrix from genotype likelihoods and used a minor allele frequency (MAF) filter of 0.05. We calculated eigen vectors and plotted the first and second principal components in R 4.1.3 using the package ggplot2 3.4.2 (Wickham 2016).

#### Mitochondrial genome assembly

We used MitoZ 3.6 (Meng et al. 2019) to create an initial *de novo* assembly of the mitochondrial genome for the *P. vaccarum* specimen from the summit of Volcán Llullaillaco, UACH8291. We then used NOVOplasty 4.3.3 (Dierckxsens et al. 2017) to generate *de novo* assemblies for the rest of samples using the mitochondrial genome for UACH8291 as a seed sequence. We annotated assembled mitochondrial genomes with MitoZ to identify coding sequences. We generated a multiple alignment of coding sequence with MAFFT 7.508 (Katoh and Standley 2013), using the --auto flag to determine the best algorithm given the data.

#### Phylogenetic analysis

Our *cytb* matrix for *Phyllotis* included newly generated sequences in combination with all non-redundant sequences available in Genbank for the species group *Phyllotis darwini* (Jayat et al. 2021; Ojeda et al. 2021) and sequences from each species in the *andium/amicus* and *osilae* species groups. As outgroups, we included sequences of the two most closely related genera in the tribe Phyllotini. We aligned sequences using Clustal W (Thompson et al. 1997), as implemented in Mega 7 (Kumar et al. 2016). We conducted phylogenetic analyses using Maximum Likelihood (ML) in IQ-TREE (Nguyen et al. 2015) using the W-IQ-TREE online implementation (http://iqtree.cibiv.univie.ac.at) (Trifinopoulos et al. 2016), with the disturbance intensity set to 0.5, the term rule set to 100, and the substitution model TPM2u+F+I+G4 found by ModelFinder (Kalyaanamoorthy et al. 2017) within IQ-TREE. Clade support was calculated via 1000 ultra-fast bootstrap pseudoreplicates.

For the whole mitochondrial genome dataset, we inferred a maximum likelihood phylogeny in IQ-TREE 2.2.0.3 (Minh et al. 2020) using 1000 ultra-fast bootstrap replicates. The substitution model used for phylogenetic inference, GTR+F+I+G4, was chosen by ModelFinder (Kalyaanamoorthy et al. 2017).

## Results and Discussion

During five high-elevation expeditions and additional low-elevation collecting trips between 2020 and 2022, we live-trapped rodents from ecologically diverse sites spanning ∼6700 m of vertical relief, from the Pacific coast of northern Chile to some of the highest summits of the Andean Cordillera (Fig. 1*A*, table S1). In the high-elevation expeditions, mammal trapping and vegetational surveys were centered on 21 volcanoes with summit elevations of 5706-6893 m (18,720-22,615 ft; fig. 1*B*, table S2). In total, we collected 498 museum voucher specimens representing 18 rodent species in 12 genera (*Abrocoma, Abrothrix, Akodon, Auliscomys, Calomys, Eligmodontia, Oligoryzomys, Phyllotis, Punomys, Mus, Rattus*, and *Octodontomys*) and two marsupial species in the genus *Thylamys* (table 1, table S3). Mice in the genera *Abrothrix* and *Phyllotis* were the most abundant taxa across the surveyed region, accounting for 27% and 49% of the total collection, respectively.

**Table 1:** Elevational ranges of capture of small mammals in the surveyed region of the Puna de Atacama (Chile/Bolivia/Argentina) and Atacama Desert of northern Chile. Maximal elevations that represent new species-specific records are shown in bold. See tables S1 and S3 for detailed locality information and capture records.

| Taxon | Elevational range of captures |
| --- | --- |
| Didelphimorphia |  |
| Didelphidae |  |
| *Thylamys elegans* | 85 m |
| *Thylamys pallidior* | 40 – 4183 m |
| Abrocomidae |  |
| *Abrocoma cinerea* | 3100 – 4543 m |
| Cricetidae |  |
| *Abrothrix andina* | 2370 – **5837 m** |
| *Abrothrix olivacea* | 4 – 3100 m |
| *Akodon albiventer* | 3380 – **5221 m** |
| *Auliscomys sp.* | 4807 m |
| *Calomys lepidus* | 4183 – 4330 m |
| *Eligmodontia hirtipes* | 4362 m |
| *Eligmodontia puerulus* | 1360 – 4099 m |
| *Oligoryzomys flavescens* | 790 m |
| *Phyllotis darwini* | 40 – 360 m |
| *Phyllotis limatus* | 650 – 3380 m |
| *Phyllotis magister* | 974 – 3380 m |
| *Phyllotis chilensis* | 3380 – **5221 m** |
| *Phyllotis vaccarum* | 49 – **6739 m** |
| *Punomys sp.* | **5461 m** |
| *Mus musculus* | 4 – 1440 m |
| *Rattus norvegicus* | 4 – 1440 m |
| Octodontidae |  |
| *Octodontomys gliroides* | 3380 m |

Vegetation surveys along the summit routes of the 21 volcanoes revealed a latitudinal trend in upper elevational limits (table S2). Among the volcanoes that we surveyed along the border of Chile and Argentina at latitudes >26° S (San Francisco, Barrancas Blancas, Vicuñas, Nevado Ojos del Salado, Los Patos, and Copiapó) (fig. 1*B*), bunch grasses in the genera *Stipa* and *Festuca* and other vascular plants typically disappeared completely at elevations between ∼4600-4900 m (table S2). By contrast, among the volcanoes that we surveyed on the border between Chile and Bolivia at latitudes <20° S (Nevado Sajama, Pomerape, Parinacota, Acotango, Guallatiri, and Sillajhuay) (fig. 1*B*), multiple species of tropical alpine herbs and dwarf shrubs exceeded elevations of 5200 m (fig. S1*A*, table S2). The flanks of Nevado Sajama harbor the highest tree line in the world, as stands of queñua (*Polylepis tarapacana*) reach elevations of ∼5200 m (Simpson 1979; fig. S1*B,C*).

Below we highlight new elevational records for multiple rodent taxa. In the case of *Phyllotis*, we report results of molecular phylogenetic and genomic analyses that confirm species identities and place new elevational records in phylogeographic context.

### Elevational Range Limits and Species Limits in Phyllotis

On the flanks of Ojos del Salado, we live-trapped six *Phyllotis* at 5250 m (tables S1, S3), 650 m above the vegetation limits on the west face of the volcano (table S2). In addition to the live-captured specimen of *P. vaccarum* from the summit of Llullaillaco (6739 m) (Storz et al. 2020), identification of active burrows of *P. vaccarum* at 6145 m on the same volcano (Steppan et al. 2022), and the discovery of desiccated cadavers (‘mummies’) of *P. vaccarum* on the summits of multiple >6000 m peaks (Storz et al. 2023), the live-capture records reported here confirm that this species is a regular denizen of the barren world of rock, ice, and snow at extreme elevations in the Puna de Atacama.

Phylogenetic analysis of *cytb* DNA sequences revealed that high-elevation *Phyllotis* specimens collected from Chilean and Bolivian Altiplano localities from Volcán Sairecabur northward (fig. 1*B*) are referrable to *P. chilensis* (*sensu* Pearson 1958)(fig. 2). By contrast, all high-elevation *Phyllotis* specimens from Altiplano localities to the south of Sairecabur fell into two distinct *cytb* clades (fig. 2), one representative of *P. vaccarum* and one that appears to represent a southern subclade of *P. limatus*. The latter finding is surprising because we collected mice with *limatus*-type *cytb* at sites in the Chilean Altiplano far south of the previously assumed range limits of *P. limatus* (fig. 3*A,B*). Previous records of mice with *limatus*-type *cytb* on the flanks of Llullaillaco were interpreted as evidence that *P. limatus* occurred at more southern latitudes (and higher elevations) than previously assumed (Storz et al. 2020). Notably, some of the highest elevation *Phyllotis* specimens that we collected south of the Tropic of Capricorn have ‘*limatus*-type’ *cytb* haplotypes, including all six mice collected from 5250 m on the flanks of Ojos del Salado, and three other mice from elevations 5070 - 6052 m on the flanks or summits of other Atacama volcanoes (figs. 2, 3*B*). If any of those specimens are indeed referable to *P. limatus*, they would far surpass the reported upper elevational range limit of that species (∼4000 m; Steppan and Ramirez 2015), and would also represent a considerable southward extension of the species’ known latitudinal range (fig. 3*A,B*). As expected the *cytb* results were corroborated by the analyses of whole mitochondrial genomes (fig. 3*C*). Since previous work identified specimens as *Phyllotis limatus* mainly on the basis of *cytb* sequences (e.g., Storz et al. 2020) and the morphological distinction among species of *Phyllotis* remains poorly understood (Ojeda et al., 2021), we attempted to obtain more conclusive insights into species limits by generating low-coverage whole-genome sequence (WGS) data for a total of 61 mice (1.26-24.1x coverage, 2.86x median). We analyzed WGS data for representative subsets of our high-elevation *Phyllotis* specimens that carried *cytb* haplotypes from the *vaccarum* clade (*n*=36) and the southern *limatus*-type subclade (*n*=15), along with specimens from more northern localities within the known range of *P. limatus* (and well outside the known range of *P. vaccarum*), all of which carried *cytb* haplotypes from the northern *limatus*-type subclade (*n*=6; fig. 3*B*). We used whole-genome sequences from *Phyllotis chilensis* (*n*=2) and *P. magister* (*n*=2) for comparisons.

**Figure 2.**
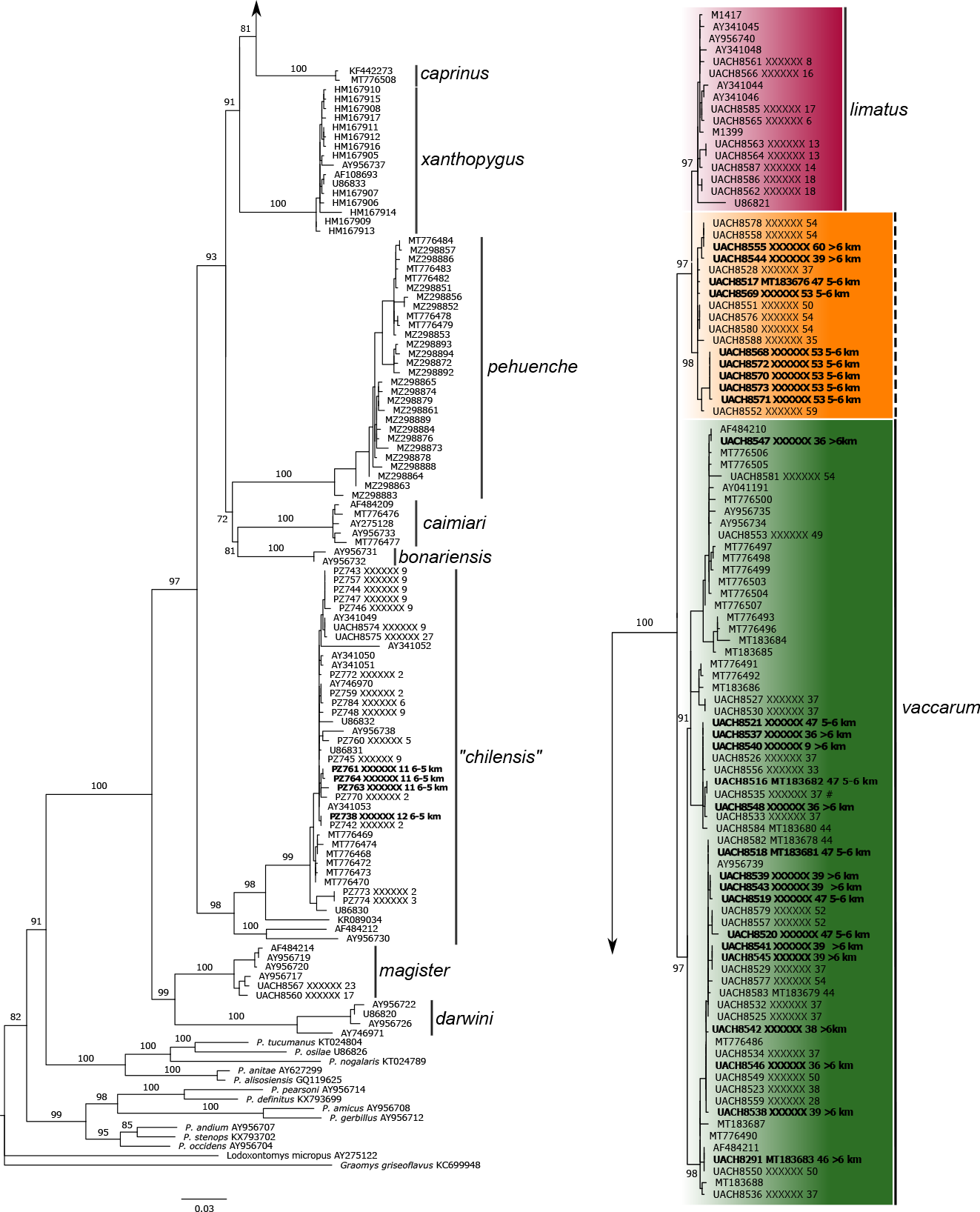
Maximum Likelihood tree (ln= -8956.735) estimated for *cytb* gene sequences of *Phyllotis*. Numbers at nodes correspond to ultra-fast bootstrap support values (only values for species and relationships among species are shown). For specimens collected as part of the present study, terminal labels refer to specimen catalog numbers, Genbank accession numbers, and localities (table S1). For sequences downloaded from GenBank, terminal labels refer to accession numbers. *Phyllotis* specimens that we collected from especially high elevations are labeled ‘5-6 km’ and ‘>6 km’. The orange clade denoted by the dashed vertical line comprises *Phyllotis* specimens with *limatus*-like *cytb* haplotypes that we collected outside the known range of *P. limatus* in northern Chile (see text for haplotypes are in fact referable to *P. vaccarum*, whereas those with the northern *limatus*-type mtDNA haplotypes are referable to *P. limatus*.

**Figure 3.**
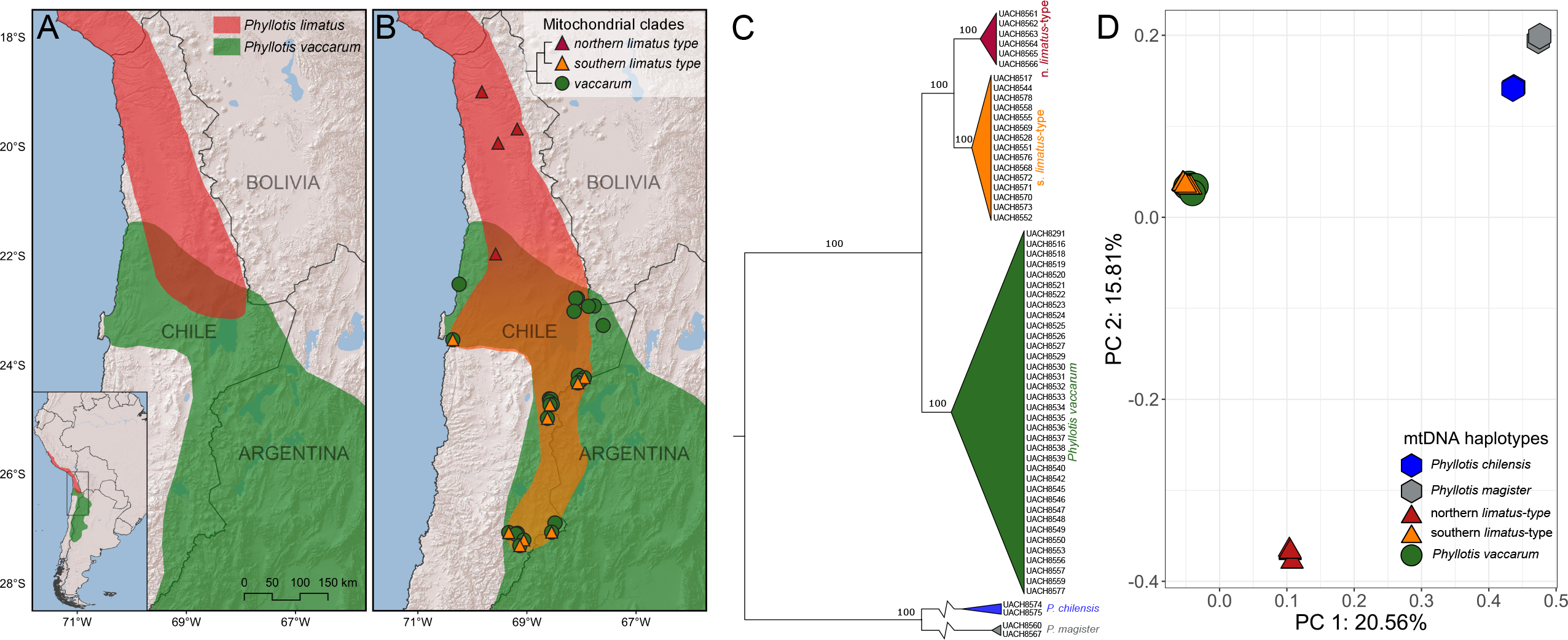
Genomic tests of alternative hypotheses regarding the latitudinal and elevational range limits of *Phyllotis limatus*. (A) Southern range limit of *Phyllotis limatus* and northern range limit of *P. vaccarum* according to traditional criteria based on morphological variation and mtDNA variation (Steppan et al. 2007; Steppan and Ramirez 2015). (B) Hypothesis for revised range limits of *P. limatus* based on the collection of mice with *limatus*-like *cytb* haplotypes in northern Chile (forming the ‘southern *limatus*’ clade shown in fig. 2) that occur well outside the traditionally assumed range of the species (but which occur well within the known range of *P. vaccarum*). Symbols denote collection localities where we recovered mice with *cytb* haplotypes that fall in three well-supported clades, one representative of *P. vaccarum*, one representative of *P. limatus* as traditionally recognized (‘northern *limatus*-type’), and one that appears to represent a southern subclade of *P. limatus* (‘southern *limatus*-type’). If mice with the ‘southern *limatus*-type’ mtDNA are in fact referable to *P. limatus*, the newly collected specimens would represent a considerable southward extension of the species’ known latitudinal range and a >1250 m upward extension of the species’ known elevational range (Steppan and Ramirez, 2015). The inset tree depicts inferred relationships among three above-mentioned *cytb* clades, according to the phylogeny estimate shown in Figure 2. (C) Estimated phylogeny based on whole mitochondrial genomes recovers the same topology as the *cytb* tree for *vaccarum* and the two *limatus* subclades (fig. 2). (D) In contrast to the relationships based on mtDNA, principal components analysis of whole-genome polymorphism data demonstrates that mice with southern *limatus*-type mtDNA group with *vaccarum* to the exclusion of mice with northern *limatus*-type mtDNA. The genomic data indicate that mice with the southern *limatus*-type mtDNA haplotypes are in fact referable to *P. vaccarum*, whereas those with the northern *limatus*-type mtDNA haplotypes are referable to *P. limatus*.

Analysis of WGS data revealed that representatives of the *vaccarum* mitochondrial clade and the southern *limatus*-type mitochondrial subclade cluster together to the exclusion of mice from the northern *limatus*-type subclade (fig. 3*D*). Results of the WGS analysis indicate that mice with the southern *limatus*-type mtDNA haplotypes are referable to *P. vaccarum*, while those with northern *limatus*-type mtDNA haplotypes represent a distinct species (*P. limatus*).

The WGS results are far easier to reconcile with the traditional understanding of the geographic distributions of the two species (fig. 3*A*) (Steppan et al. 2007; Steppan and Ramirez 2015). The WGS data also confirm that all extreme high-elevation *Phyllotis* mice from elevations 5070-6739 m on the flanks or summits of five different volcanoes (Púlar, Salín, Llullaillaco, Ojos del Salado, and Copiapó) are referable to *P. vaccarum*, not *P. limatus*. Causes of the discordance between mitochondrial and nuclear genomic variation in these mice remains to be elucidated, and could potentially reflect incomplete lineage sorting and/or a history of introgressive hybridization between *P. limatus* and *P. vaccarum*. Whatever the cause, the results demonstrate that mtDNA sequence data alone do not always provide a reliable basis for distinguishing species of *Phyllotis*.

Notably, high-elevation specimens of *Phyllotis vaccarum* exhibited close genetic affinities with multiple specimens that we collected at or near sea level along the coast of northern Chile (fig. 2). Thus, this species not only inhabits higher elevations than any other mammal, it also has by far the broadest elevational range, from sea-level to >6700 m.

### Elevational Records Provide a New Appreciation of Environmental Limits of Vertebrate Life

Our high-elevation surveys yielded new elevational records for five rodent species, including previously reported records for *Phyllotis vaccarum* (Storz et al. 2020) and *Punomys lemminus* (Quiroga-Carmona et al. 2023) (table 1). Our record capture of *Abrothrix andina* at 5837 m on the flanks of Ojos del Salado far surpasses the previously reported limit of ∼5000 m (Mann 1978; Patterson et al. 2015). In western Bolivia, we captured multiple *Akodon albiventer* from 5020-5221 m on the flanks of Nevado Sajama and Volcán Parinacota (the species had not been previously recorded >4500 m; Pardiñas et al. 2015) and we captured multiple *Phyllotis chilensis* from 5027-5221 m on the flanks of Acotango and Parinacota (the species had not been previously recorded >4700 m; Rengifo et al. 2022). We use the name ‘*chilensis*’, following Pearson (1958), but we note that the taxonomy of *Phyllotis* in the Altiplano is in a state of flux, and names could change in the future.

Prior to our Andean surveys, the mammalian species with the highest specimen-based elevational record was the long-eared pika, *Ochotona macrotis*, with two Himalayan specimens from 5182 m (US National Museum 198648 and 198649). In the Andean Altiplano, the highest records reported by Rengifo et al. (2022) were for *Abrothrix jelskii* from 5100 m in the Central Andean puna of southern Peru (Museo de Historia Natural, Universidad Mayor de San Marcos, MUSM37711-37727). Our high-elevation Andean expeditions yielded a total of 26 specimens of five different species that surpass those elevations, including the highest elevational record for mammals (*Phyllotis vaccarum*) as well as species with the 2^nd^ through 5^th^ highest records (table 1). These specimen-based records greatly extend the known limits of what can be considered habitable environments for mammals.

### New Frontiers of Biological Surveys

Biological surveys of the abyssal and hadal zones of deep ocean trenches have yielded novel discoveries about the environmental limits of marine life by using specialized approaches (e.g., manned and unmanned submersibles) distinct from those employed in surveys of upper pelagic zones. Likewise, surveys of terrestrial mammals at extreme elevations require the specialized logistics and acclimatization protocols of a full-fledged mountaineering expedition, with the establishment of base camps and high camps for the daily monitoring of trap lines. Using this approach, our trapping surveys of Nevado Sajama, Volcán Llullaillaco, Ojos del Salado, and other >6000 m volcanoes have yielded surprising elevational records for multiple rodent species, all of which are cataloged as voucher specimens. These records provide valuable baseline data for monitoring effects of climate change on the elevational distributions of Andean mammals, and they also prompt new questions about basic ecology and mechanisms of physiological adaptation to environmental extremes.

## Acknowledgments

This work was funded by grants from the National Institutes of Health (R01 HL159061, JFS and JMG), National Science Foundation (IOS-2114465, JFS; OIA-1736249, JFS and JMG), National Geographic Society (NGS-68495R-20, JFS), and the Fondo Nacional de Desarrollo Científico y Tecnológico (Fondecyt 1221115, GD). We thank Mario Pérez-Mamani, Juan Carlos Briceño, Luís Castillo, and Alex Damian González Sandoval for assistance and companionship in the field.

## Statement of Authorship

J.F.S., S.L., M.Q-C., G.D., and J.M.G designed the research; J.F.S, M.Q-C., N.M.B, J. S-B., A.R.C. and G.D. performed the field work; S.L. performed the laboratory work; S.L., M.Q-C., J. S-B., J.C.O., G.D., and J.M.G. analyzed data; J.F.S., S.L., M.Q-C., N.M.B., G.D., and J.M.G. prepared figures and wrote the manuscript.

## Data and Code Availability

All sequence data reported in this study are archived in the National Center for Biotechnology Information/GenBank. Raw sequencing reads are available in the Sequence Read Archive, and all genome sequences were deposited in GenBank under BioProject PRJNA950396 and are publicly available as of the date of publication.

