## Supplementary Material for "Extreme high-elevation mammal surveys reveal unexpectedly high upper range limits of Andean mice"

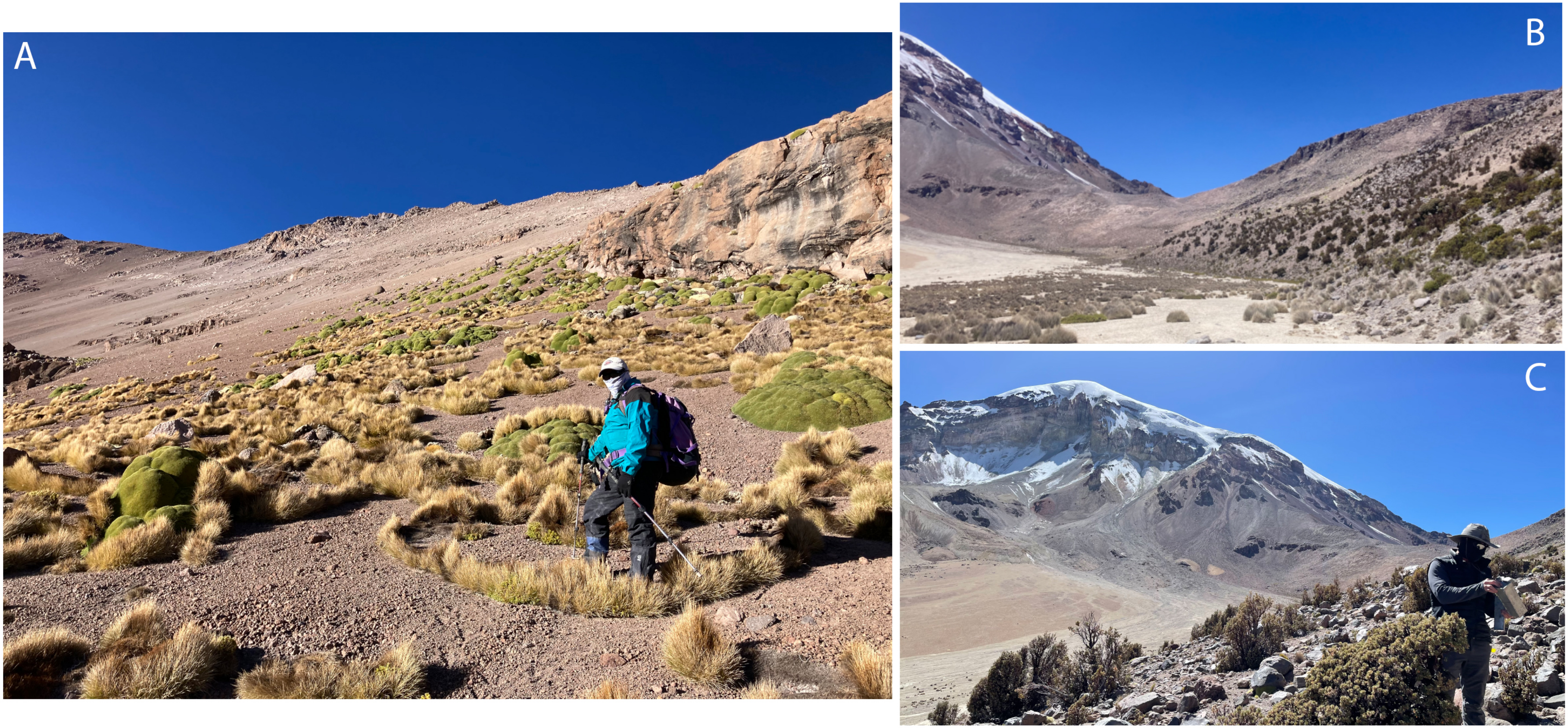


**Figure S1:** Examples of upper elevational limits of vegetation. (A) View of vegetation limits at ~5300 m on the flanks of Volcán Sillajhuay, Bolivia-Chile (Departamento de Oruro/Región de Arica y Parinacota), 19°44′32″S, 68°41′26″W. (B) Example of the highest tree line in the world: stands of queñua (*Polylepis tarapacana*) at ~5100 m on the flanks of Volcán Nevado Sajama, Bolivia (Departamento de Oruro), 18°06′29"S, 68°52′59″W. (C) Placement of Sherman traps along queñua tree line at 4880 m on the flanks of Volcán Nevado Sajama, Bolivia (Departamento de Oruro), 18°06.77′S, 68°54.93′W.

**Table S1**. Trapping localities in northern Chile and bordering regions of Argentina and Bolivia.

| n° | Locality description | Elevation | Coordinates |
|---|---|---|---|
| 1. | Chile, Arica y Parinacota, campamento Laguna Casiri | 4807 m | 18°04.25′S, 69°04.90′W |
| 2. | Bolivia, Oruro, Geiser Sajama | 4420 m | 18°05.77′S, 69°01.93′W |
| 3. | Bolivia, Oruro, Geiser Sajama | 4362 m | 18°06.34′S, 69°00.97′W |
| 4. | Bolivia, Oruro, 1 Km N de campamento base, campo de rocas | 5020 m | 18°06.35′S, 68°54.55′W |
| 5. | Bolivia, Oruro, Geiser Sajama | 4330 m | 18°06.61′S, 69°00.67′W |
| 6. | Bolivia, Oruro, Volcán Sajama, campamento base, | 4880 m | 18°06.77′S, 68°54.93′W |
| 7. | Bolivia, Oruro, Camino Sajama | 4290 m | 18°06.89′S, 68°58.02′W |
| 8. | Bolivia, Oruro, Camino Sajama | 4295 m | 18°07.08′S, 68°58.04′W |
| 9. | Bolivia, Oruro, Sajama, mirador Monte Cielo | 4537 m | 18°08.13′S, 68°57.28′W |
| 10. | Bolivia, Oruro, Sajama, mitad del camino al mirador Monte Cielo | 4377 m | 18°08.15′S, 68°57.57′W |
| 11. | Bolivia, Oruro, Volcán Parinacota, campamento alto | 5221 m | 18°09.13′S, 69°07.34′W |
| 12. | Bolivia, Oruro, Volcán Acotango, campamento base | 5027 m | 18°21.98′S, 69°01.45′W |
| 13. | Chile, Arica y Parinacota, Camarones, Quebrada de Camarones, Ruta A-345, km 28 | 790 m | 19°00.17′S, 69°49.62′W |
| 14. | Chile, Arica y Parinacota, Camarones, Quebrada de Camarones, Ruta A-345, km 20.7 | 650 m | 19°00.59′S, 69°53.65′W |
| 15. | Chile, Arica y Parinacota, Laguna Cota Kulco | 4183 m | 19°38.62′S, 68°45.58′W |
| 16. | Chile, Arica y Parinacota, Huara, Chusmiza, Quebrada de Ocharaza | 3380 m | 19°41.00′S, 69°10.80′W |
| 17. | Chile, Tarapacá, Huara, Quebrada de Tarapacá, Quillahuasa | 1440 m | 19°54.93′S, 69°29.77′W |
| 18. | Chile, Tarapacá, Huara, Quebrada de Tarapacá, Huarasiña | 1360 m | 19°56.48′S, 69°31.85′W |
| 19. | Chile, Antofagasta, Ollagüe, Volcán Aucanquilcha | 4543 m | 21°11.26′S, 68°24.24′W |
| 20. | Chile, Antofagasta, Ollagüe, Volcán Aucanquilcha | 4510 m | 21°11.26′S, 68°24.10′W |
| 21. | Chile, Antofagasta, Tocopilla, Desembocadura del Rio Loa | 15 m | 21°25.55′S, 70°02.58′W |
| 22. | Chile, Antofagasta, Tocopilla, Desembocadura del Rio Loa | 4 m | 21°25.68′S, 70°03.14′W |
| 23. | Chile, Antofagasta, María Elena, Rio Loa | 974 m | 21°57.34′S, 69°33.81′W |
| 24. | Chile, Antofagasta, María Elena, Rio Loa, Paso Toco (Puente Teresa) | 1100 m | 21°58.14′S, 69°34.44′W |
| 25. | Chile, Antofagasta, Tocopilla, Cobija | 49 m | 22°31.32′S, 70°14.65′W |
| 26. | Chile, Antofagasta, Ollagüe, Cerro Colorado, campamento base | 4840 m | 22°35.05′S, 67°54.32′W |
| 27. | Chile, Antofagasta, San Pedro de Atacama, base del Cerro Sairecábur | 4480 m | 22°43.63′S, 67°53.36′W |
| 28. | Chile, Antofagasta, San Pedro de Atacama, Ruta 27 CH km 33 | 4099 m | 22°46.69′S, 68°04.62′W |
| 29. | Chile, Antofagasta, San Pedro de Atacama, Gachi, Sector Guatín, Margen Río Puritana | 3120 m | 22°46.89′S, 68°06.33′W |
| 30. | Chile, Antofagasta, San Pedro de Atacama, Ruta 27 CH km 45.5 | 4750 m | 22°55.20′S, 67°46.04′W |
| 31. | Chile, Antofagasta, San Pedro de Atacama, Ruta B-245 km 20 | 3240 m | 22°55.54′S, 67°52.38′W |
| 32. | Chile, Antofagasta, San Pedro de Atacama, Ruta B-241 km 117 | 2370 m | 23°01.09′S, 68°08.18′W |
| 33. | Chile, Antofagasta, San Pedro de Atacama, Volcán Acamarachi, campamento base | 4895 m | 23°16.57′S, 67°36.21′W |
| 34. | Chile, Antofagasta, San Pedro de Atacama, caldera de Volcán Acamarachi | 5461 m | 23°16.98′S, 67°37.46′W |
| 35. | Chile, Antofagasta, Antofagasta, Reserva Natural La Chimba, Quebrada La Chimba | 420 m | 23°32.27′S, 70°21.55′W |

| | | | |
|---|---|---|---|
| 36. | Chile, Antofagasta, San Pedro de Atacama, cumbre de Volcán Púlar | 6233 m | 24°11.52′S, 68°03.38′W |
| 37. | Chile, Antofagasta, San Pedro de Atacama, Salar de Púlar, Volcán Púlar | 3651 m | 24°13.97′S, 67°56.88′W |
| 38. | Chile, Antofagasta, San Pedro de Atacama, Salar de Púlar, Volcán Salín | 4016 m | 24°16.84′S, 68°00.68′W |
| 39. | Chile (Antofagasta, San Pedro de Atacama), cumbre de Volcán Salín | 6029 m | 24°19.70′S, 68°04.04′W |
| 40. | Chile, Antofagasta, Antofagasta. Volcán Socompa | 4555 m | 24°25.17′S, 68°15.91′W |
| 41. | Chile, Taltal, Paposo, Ruta 1, km 90 | 85 m | 24°31.00′S, 70°31.00′W |
| 42. | Chile, Antofagasta, Antofagasta, Parque Nacional Llullaillaco, Refugio Aguadas de Zorritas | 4150 m | 24°37.16′S, 68°35.33′W |
| 43. | Chile, Antofagasta, Antofagasta, Parque Nacional Llullaillaco, 300 m al norte del Refugio Aguadas de Zorritas | 4140 m | 24°37.29′S, 68°35.22′W |
| 44. | Chile, Antofagasta, Antofagasta, Parque Nacional Llullaillaco, 3 km al suroeste del Refugio Aguadas de Zorritas | 4360 m | 24°37.74′S, 68°33.58′W |
| 45. | Chile, Antofagasta, Antofagasta, Parque Nacional Llullaillaco, Volcán Llullaillaco, campamento base – Ruta Norte | 4620 m | 24°40.53′S, 68°34.84′W |
| 46. | Chile (Antofagasta, Antofagasta), cumbre del Volcán Llullaillaco | 6739 m | 24°43.21′S, 68°32.22′W |
| 47. | Chile, Antofagasta, Antofagasta, Parque Nacional Llullaillaco, Volcán Llullaillaco, campamento base – Ruta Este | 5070 m | 24°43.81′S, 68°34.68′W |
| 48. | Chile, Antofagasta, Taltal, Paposo, Ruta 1, km 95.7 | 40 m | 24°51.18′S, 70°31.40′W |
| 49. | Chile, Antofagasta, Taltal, Paposo, Ruta 1, km 85.7 | 45 m | 24°55.77′S, 70°30.77′W |
| 50. | Chile, Antofagasta, Antofagasta, Salar de Aguas Calientes | 3750 m | 24°58.56′S, 68°37.53′W |
| 51. | Chile, Antofagasta, Taltal, Quebrada Cachina, Ruta B 980 | 360 m | 25°54.03′S, 70°36.67′W |
| 52. | Chile, Atacama, Copiapó, Refugio Laguna Verde | 4460 m | 26°53.42′S, 68°29.16′W |
| 53. | Chile, Atacama, Copiapó, Ojos del Salado, campamento base, Refugio Atacama | 5250 m | 27°03.59′S, 68°32.86′W |
| 54. | Chile, Atacama, Copiapó, Vallecitos, Pajas Grandes (Colas de Zorro) | 3100 m | 27°03.97′S, 69°20.04′W |
| 55. | Chile, Atacama, Copiapó, Parque Nacional Nevado de Tres Cruces, Refugio Maricunga | 3780 m | 27°04.84′S, 69°10.54′W |
| 56. | Chile, Atacama, Copiapó, campamento alto, Ojos del Salado | 5837 m | 27°05.25′S, 68°32.29′W |
| 57. | Chile, Atacama, Copiapó, Parque Nacional Nevado de Tres Cruces, roquerio frente a Laguna Santa Rosa, cenizas volcánica | 3780 m | 27°05.42′S, 69°10.49′W |
| 58. | Chile, Atacama, Copiapó, Parque Nacional Nevado de Tres Cruces, Ruinas Incaicas, cerca de Laguna Santa Rosa | 3868 m | 27°05.77′S, 69°12.21′W |
| 59. | Chile, Atacama, Copiapó, campamento base, Volcán Copiapó | 4100 m | 27°12.21′S, 69°03.47′W |
| 60. | Chile, Atacama, Copiapó, cumbre de Volcán Copiapó | 6052 m | 27°18.35′S, 69°07.85′W |

**Table S2**. Surveyed volcanoes in the Puna de Atacama of northern Chile and bordering regions of Argentina and Bolivia.

| | Volcano | Elevation | Vegetational limit[1] | Coordinates (summit) |
|---|---|---|---|---|
| 1. | Nevado Sajama, Bolivia, Departamento de Oruro | 6542 m | 5200 m | 18°06′29″S, 68°52′59″W |
| 2. | Volcán Pomerape, Boliva-Chile (Departamento de Oruro/Región de Arica y Parinacota) | 6282 m | 5500 m | 18°07′33″S, 69°07′39″W |
| 3. | Volcán Parinacota, Boliva-Chile (Departamento de Oruro/Región de Arica y Parinacota) | 6319 m | 5400 m | 18°09′57″S, 69°08′30″W |
| 4. | Acotango, Bolivia-Chile (Departamento de Oruro/Región de Arica y Parinacota) | 6052 m | 5200 m | 18°22′56″S, 69°02′52″W |
| 5. | Volcán Guallatiri, Chile (Región de Arica y Parinacota) | 6071 m | 5300 m | 18°25′00″S, 69°05′30″W |
| 6. | Volcán Sillajhuay, Bolivia-Chile (Departamento de Oruro/Región de Arica y Parinacota) | 5982 m | 5300 m | 19°44′32″S, 68°41′26″W |
| 7. | Volcán Aucanquilcha, Chile (Región de Antofagasta) | 6176 m | 4900 m | 21°13′00″S,68°28′00″W |
| 8. | Cerro Colorado, Chile (Región de Antofagasta) | 5748 m | 4800 m | 22°35.54′S, 67°55.33′W |
| 9. | Cerro Sairecabur, Chile (Región de Antofagasta) | 5971 m | 4600 m | 22°43′00″S, 67°53′30″W |
| 10. | Volcán Acamarachi, Chile (Región de Antofagasta) | 6046 m | 5200 m | 23°17′45″S, 67°37′02″W |
| 11. | Volcán Púlar, Chile (Región de Antofagasta) | 6233 m | 4700 m | 24°12′02″S, 68°03′59″W |
| 12. | Volcán Salín, Argentina-Chile (Provincia de Salta/Región de Antofagasta) | 6029 m | 4600 m | 24°19′54″S, 68°03′50″W |
| 13. | Volcán Socompa, Argentina-Chile (Provincia de Salta/Región de Antofagasta) | 6051 m | 4400 m | 24°23′36″S, 68°15′22″W |
| 14. | Volcán Llullaillaco, Argentina-Chile (Provincia de Salta/Región de Antofagasta) | 6739 m | 5100 m | 24°43′12″S, 68°32′13″W |
| 15. | Volcán Lastarria, Argentina-Chile (Provincia de Salta/Región de Antofagasta) | 5706 m | 4800 m | 25°09′27″S, 68°30′34″W |
| 16. | Volcán San Francisco, Argentina-Chile (Provincia de Catamarca/Región de Atacama) | 6016 m | 4900 m | 26°55′06″S68°15′47″W |
| 17. | Volcán Barrancas Blancas, Chile (Región de Atacama) | 6119 m | 4600 m | 26°59'28"S, 68°39'58"W |
| 18. | Cerro Vicuñas, Chile (Región de Atacama) | 6067 m | 4700 m | 27°01′17″S, 68°37′08″W |
| 19. | Nevado Ojos del Salado, Argentina-Chile (Provincia de Catamarca/Región de Atacama) | 6893 m | 4600 m | 27°06′34″S, 68°32′32″W |
| 20. | Volcán Los Patos, Argentina-Chile (Provincia de Catamarca/Región de Atacama) | 6239 m | 4700 m | 27°18′04″S, 68°48′32″W |
| 21. | Volcán Copiapó, Chile (Región de Atacama) | 6052 m | 4700 m | 27°18′21″S, 69°07′51″W |

[1]Rounded to nearest 100 m above sea level (see *Material and Methods* for survey protocol).

**Table S3. Number of captures per species, per locality**. See table S1 for geographic coordinates and other details for all trapping localities.

| | Locality | | | | | | | | | | | | | | | | | | | | | | | | | | | | | |
|---|---|---|---|---|---|---|---|---|---|---|---|---|---|---|---|---|---|---|---|---|---|---|---|---|---|---|---|---|---|---|
| **Species** | 1 | 2 | 3 | 4 | 5 | 6 | 7 | 8 | 9 | 10 | 11 | 12 | 13 | 14 | 15 | 16 | 17 | 18 | 19 | 20 | 21 | 22 | 23 | 24 | 25 | 26 | 27 | 28 | 29 | 30 |
| *Thylamys elegans* | | | | | | | | | | | | | | | | | | | | | | | | | | | | | | |
| *Thylamys pallidior* | | | | | | | | | | | | | X | | X | X | | | | | | | | | | | | | | |
| *Abrocoma cinerea* | | | | | | | | | | | | | | | | | | | X | | | | | | | | | | | |
| *Abrothrix andina* | X | | | | | | | | | | | | | | | | | | | | | | | | | | | X | | |
| *Abrothrix olivacea* | | | | | | | | | | | | | X | X | | | X | X | | | X | X | X | X | | | | | | |
| *Akodon albiventer* | | X | X | X | X | X | X | X | X | | X | | | | X | X | | | | | | | | | | | | | | |
| *Auliscomys sp.* | X | | | | | | | | | | | | | | | | | | | | | | | | | | | | | |
| *Calomys lepidus* | | | | | X | | | | | | | | | | X | | | | | | | | | | | | | | | |
| *Eligmodontia hiritipes* | | | X | | | | | | | | | | | | | | | | | | | | | | | | | | | |
| *Eligmodontia puerulus* | | | | | | | | | | | | | | | | | | X | | | | | | | | | | X | | |
| *Oligoryzomys flavescens* | | | | | | | | | | | | | X | | | | | | | | | | | | | | | | | |
| *Phyllotis darwini* | | | | | | | | | | | | | | | | | | | | | | | | | | | | | | |
| *Phyllotis limatus* | | | | | | | | | | | | | X | X | | X | X | X | | | | | X | | | | | | | |
| *Phyllotis magister* | | | | | | | | | | | | | | | | X | X | | | | | | X | X | | | | | | |
| *Phyllotis chilensis* | X | X | X | | X | X | | | X | X | X | X | | | X | | | | X | X | | | | | | X | X | | | |
| *Phyllotis vaccarum* | | | | | | | | | | | | | | | | | | | | | | | | | X | | | X | X | X |
| *Punomys sp.* | | | | | | | | | | | | | | | | | | | | | | | | | | | | | | |
| *Mus musculus* | | | | | | | | | | | | | X | | | | X | X | | | X | X | X | X | | | | | | |
| *Rattus norvegicus* | | | | | | | | | | | | | X | | | | X | | | | X | X | | X | | | | | | |
| *Octodontomys gliroides* | | | | | | | | | | | | | | | | X | | | | | | | | | | | | | | |

| | Locality | | | | | | | | | | | | | | | | | | | | | | | | | | | | | |
|---|---|---|---|---|---|---|---|---|---|---|---|---|---|---|---|---|---|---|---|---|---|---|---|---|---|---|---|---|---|---|
| **Species** | 31 | 32 | 33 | 34 | 35 | 36 | 37 | 38 | 39 | 40 | 41 | 42 | 43 | 44 | 45 | 46 | 47 | 48 | 49 | 50 | 51 | 52 | 53 | 54 | 55 | 56 | 57 | 58 | 59 | 60 |
| *Thylamys elegans* | | | | | | | | | | | X | | | | | | | | | | | | | | | | | | | |
| *Thylamys pallidior* | | X | | | | | | | | | | | | | | | | X | | | | | | | | | | | | |
| *Abrocoma cinerea* | | | | | | | | | | | | | | | | | | | | | | | | X | | | | | | |
| *Abrothrix andina* | | X | | | | | X | | | X | | X | X | X | X | | | | | X | | X | | X | | X | | X | | |
| *Abrothrix olivacea* | | | | | | | | | | | | | | | | | | X | X | | X | | | X | | | | | | |
| *Akodon albiventer* | | | | | | | | | | | | | | | | | | | | | | | | | | | | | | |
| *Auliscomys sp.* | | | | | | | | | | | | | | | | | | | | | | | | | | | | | | |
| *Calomys lepidus* | | | | | | | | | | | | | | | | | | | | | | | | | | | | | | |
| *Eligmodontia hiritipes* | | | | | | | | | | | | | | | | | | | | | | | | | | | | | | |
| *Eligmodontia puerulus* | | | | | | | | | | | | | | | | | | | | | | | | | | | | | | |
| *Oligoryzomys flavescens* | | | | | | | | | | | | | | | | | | | | | | | | | | | | | | |
| *Phyllotis darwini* | | | | | | | | | | | | | | | | | | X | X | | X | | | | | | | | | |
| *Phyllotis limatus* | | | | | | | | | | | | | | | | | | | | | | | | | | | | | | |
| *Phyllotis magister* | | | | | | | | | | | | | | | | | | | | | | | | | | | | | | |
| *Phyllotis chilensis* | | | | | | | | | | | | | | | | | | | | | | | | | | | | | | |
| *Phyllotis vaccarum* | X | X | X | | X | X | X | X | X | | | X | | X | X | X | X | | | X | | X | X | X | X | | X | X | X | X |
| *Punomys sp.* | | | | X | | | | | | | | | | | | | | | | | | | | | | | | | | |
| *Mus musculus* | | | | | | | | | | | | | | | | | | | | | | | | | | | | | | |
| *Rattus norvegicus* | | | | | | | | | | | | | | | | | | | | | | | | | | | | | | |
| *Octodontomys gliroides* | | | | | | | | | | | | | | | | | | | | | | | | | | | | | | |
